# Inferring a directed acyclic graph of phenotypes from GWAS summary statistics

**DOI:** 10.1101/2023.02.10.528092

**Authors:** Rachel Zilinskas, Chunlin Li, Xiaotong Shen, Wei Pan, Tianzhong Yang

## Abstract

Estimating phenotype networks is a growing field in computational biology. It deepens the understanding of disease etiology and is useful in many applications. In this study, we present a method that constructs a phenotype network by assuming a Gaussian linear structure model embedding a directed acyclic graph (DAG). We utilize genetic variants as instrumental variables and show how our method only requires access to summary statistics from a genome-wide association study (GWAS) and a reference panel of genotype data. Besides estimation, a distinct feature of the method is its summary statistics-based likelihood ratio test on directed edges. We applied our method to estimate a causal network of 29 cardiovascular-related proteins and linked the estimated network to Alzheimer’s disease (AD). A simulation study was conducted to demonstrate the effectiveness of this method. An R package sumdag implementing the proposed method, all relevant code, and a Shiny application are available at https://github.com/chunlinli/sumdag.

## 1. Introduction

Network analysis has deepened our understanding of biological mechanisms and disease etiologies (Zhang and Itan, 2019). Specifically, protein-protein interaction (PPI) networks that capture the interplay of proteins in the biomolecular systems are vital for normal cell functions (Snider et al., 2015). Disturbing of the normal pattern in the PPI network can be causative to or indicative of a disease state. Studies have linked co-regulatory networks of proteins to a variety of complex diseases (Ross and Poirier, 2004; Emilsson et al., 2018). Recently, a network-based method modeling PPI boasted high accuracy rates in cancer prediction (Id et al., 2021). Cheng et al. (2021) further showed that disease-associated variants were significantly enriched in the sequences coding PPI interfaces compared to variants in healthy individuals. Their work also demonstrated associations of PPIs with drug resistance and overall survival, highlighting the use of protein networks for informing genotype-based therapy. Network-based analyses have shown their potential in advancing precision medicine for complex diseases over traditional approaches which focus on monogenic mutations and independent assessment of risk factors (Napoli et al., 2020).

Network analyses can be categorized into two groups. One utilizes only phenotypic data to construct networks. For example, weighted gene network co-expression analysis estimates an undirected network which is further characterized using dimension reduction techniques (Zhang and Horvath, 2005). Graphical lasso formulation employs penalized methods to estimate a Gaussian graphical model for a large number of variables (Witten et al., 2012). Bayesian network analysis (Friedman et al., 2000) estimates directed acyclic graphs (DAG), which are widely accepted in biological systems (Ashburner et al., 2000), and its recent improvements in computational approaches have led to much shortened computational time (Liu et al., 2016). The other group of methods exploits the use of instrumental variable (IV) techniques to estimate a DAG, assuming a linear structural equation model. Chen et al. (2018) developed a penalized two-stage least squares method to estimate a DAG, assuming known intervention targets. Li et al. (2023) further extended the work to accommodate unknown intervention targets commonly encountered in biological applications.

Individual-level data are required for all the methods above, which, however, can be difficult to obtain, especially for human studies, due to logistic limitations and privacy concerns. On the other hand, many genome-wide association studies (GWAS) have shared their summary statistics publicly, generating a rich and valuable data resource. Thus, we propose adapting the network estimation and inference methods of Li et al. (2023) to rely only on GWAS summary statistics and a genetic reference panel, both much more easily accessible. We will show how a DAG can be estimated for cardiovascular-related proteins using a large-scale proteomic GWAS summary dataset, and then link the protein network to Alzheimer’s disease (AD). The algorithm for the proposed work is packaged in R. Our work represents one of the initial attempts to utilize GWAS summary statistics in the construction of a DAG. We expect that our work can facilitate more comprehensive network analysis in studying biological and medical relationships. In addition to inferring PPI network, our method is readily applicable to understand the interplay of many other molecular and non-molecular phenotypes, as long as the corresponding GWAS summary statistics are available.

## 2. Methods

### 2.1. Network modeling and data

#### 2.1.1. Directed phenotype network

Our goal is to use genotypes as external interventions to construct and infer a DAG that describes the directed relationships among a set of phenotypes. In the framework of interventional Gaussian DAG (Li et al., 2023), we assume

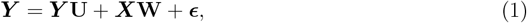

where ***Y*** = (***y***_1_, …, ***y***_*P*_ ) is the *N × P* data matrix of *P* phenotypes, ***X*** = (***x***_1_, …, ***x***_*Q*_) is the *N × Q* data matrix of *Q* genotypes serving as IVs, ***ϵ*** = (***ϵ***_1_, …, ***ϵ***_*P*_ ) is the *N × P* error matrix with each row sampled from 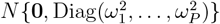, and *N* is the sample size. Note that Equation (1) lacks an intercept because we assume phenotype and genotype are centered at mean 0, which could be easily done with individual-level GWAS data.

In Equation (1), **U** and **W** are unknown parameters to be estimated. The *P × P* matrix **U** = (u_*kj*_) specifies the network structure such that u_*kj*_≠ 0 indicates a directed relation from phenotype *k* to phenotype *j*. The *Q × P* matrix **W** = (w_*qp*_) specifies the targets and strengths of interventions in that w_*qp*_≠ 0 indicates an interventional relation from genotype *q* to phenotype *p*. Let *ℰ* = *{*(*k, j*) : u_*kj*_≠ 0} be the set of directed relations and ℐ = *{*(*q, p*) : w_*qp*_≠ 0} be the set of interventional relations.

#### 2.1.2. Summary statistics and reference panel

In Equation (1), the data matrices ***X*** and ***Y*** contain individual information such that each row represents the variables measured on an individual. On the other hand, GWAS summary statistics aggregate *N* observations into a single measure for each single nucleotide polymorphism (SNP) across the whole genome. This measure is the average effect of having one copy of the effect allele of the SNP on the phenotype being studied. It is estimated by 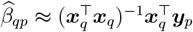, often reported along with accompanying statistics such as the corresponding standard error 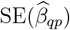, z-score *z*_*qp*_, sample size *N*, reference allele (REF), minor allele frequency (MAF) and p-value. The summary statistics of the *Q* SNPs in **W** are included in the GWAS summary data.

As a complement to the summary-level data, a reference panel comprising genotypic data of individuals from a general population provides the correlation structure among the genotypes. Many existing resources can be used for such a reference panel (The 1000 Genomes Project Consortium, 2015; International HapMap Consortium, 2005; Taliun et al., 2021; Bycroft et al., 2018). Given an *N*_*r*_ *× Q* (centered) reference panel ***X***_*r*_ of *N*_*r*_ individuals, we follow the conventional suggestion (Mak et al., 2017) to regularize the genetic correlation matrix ***R*** such that 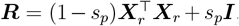 where 0 ⩽ *s*_*p*_ ⩽ 1 is a real number controlling the degree of regularization.

From the summary statistics and the reference panel, we compute the following quantities that are used for the construction and inference of the directed phenotype network. The subsequent computation also assumes ***X*** and ***Y*** are centered, which does not influence 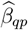 and the accompanying statistics.

- The covariance matrix of genotypes 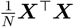 is estimated by 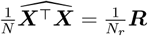
- Let 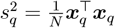. Then 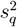 is estimated by 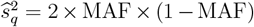 provided that MAF is reported in the summary statistics, or otherwise estimated by 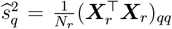 the *q*^*th*^ diagonal element of 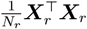.
- Given 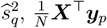 is estimated by 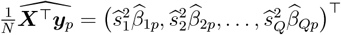
- For 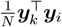, we use the median estimate 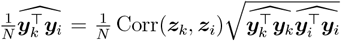.
- Finally, 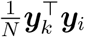 is estimated using the null SNPs from GWAS summary statistics, i.e., SNPs not marginally associated with ***y***_*k*_ or ***y***_*i*_. Following Kim et al. (2015), Corr(***y***_*k*_, ***y***_*i*_) ≈ Corr(***z***_*k*_, ***z***_*i*_) where ***z***_*k*_ and ***z***_*i*_ are vectors of z-scores for the null SNPs. Thus, we can rearrange the sample correlation formula for (centered) phenotype variables and plug in our approximation to obtain 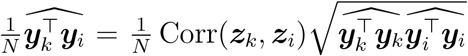 In practice, the SNPs with p-values larger than 0.05 are considered as null SNPs. An alternative method to consider for estimating 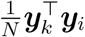 is also feasible to use for GWAS (Bulik-Sullivan et al., 2015), although not used herein.

Next we extend the framework of interventional Gaussian DAG to leverage large-scale GWAS summary statistics.

### 2.2. Method for network construction

The estimation of interventional Gaussian DAG consists of three steps.

(E1) First, we use penalized regressions to estimate the genotype-phenotype association matrix **V** := **W**(**I** *−* **U**)^*−*1^ in the following equation

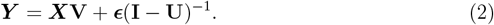

(E2) Next, we employ the peeling algorithm (Li et al., 2023) to learn a super-DAG, i.e., a directed super-graph without cycles, based on 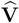 obtained in Step (E1).

(E3) Finally, we estimate **U** and **W** through penalized regressions based on the estimated super-DAG in Step (E2).

Now, we elaborate on our extensions to accommodate summary statistics.

#### 2.2.1. Estimation of **V** by truncated Lasso penalized regressions

In Equation (2), the matrix **V** can be estimated column-wise from

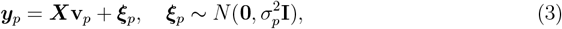

where ***y***_*p*_ is the data vector of phenotype *p*, ***X*** is the data matrix of genotypes, and vector **v**_*p*_ = (v_1*p*_, …, v_*Qp*_)^*T*^ is the *p*^*th*^ column of **V**. Given the summary statistics, we expand the squared error function 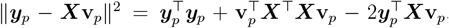, and replace the quantities 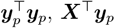, and ***X***^*T*^***X*** in Equation (3) with their estimates 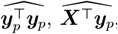, and 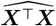, respectively. As a result, we estimate **v**_*p*_ through regressions with the Truncated Lasso Penalty (TLP) (Shen et al., 2012) to minimize

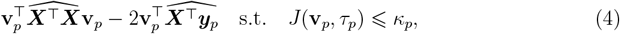

where *κ*_*p*_ *>* 0 is an integer tuning parameter and 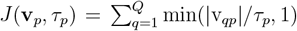 is the TLP function, which does not penalize the parameters over the threshold *τ*_*p*_. We use the R package “glmtlp” (Li et al., 2022) to fit the summary-level data regression (4).

For implementation, we fix 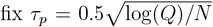 and choose *κ*_*p*_ ∈ {1, …, *Q*} individually for each of the *P* penalized regressions by minimizing the pseudo-BIC (Pattee and Pan, 2020), which is defined as 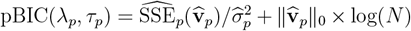, where 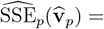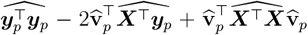 is the (estimated) sum of squared error of 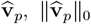 is the number of nonzero coefficients in 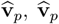 is the estimate in (4) with tuning parameters (λ_p_, τ _p_)*N* is the sample size (when *N* differs, the median is taken), and 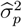 is the estimated residual variance for phenotype *p* in Equation (3). When *Q* is small compared to *N* as in our application, a consistent estimate for 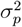 can be obtained from the ordinary least squares using all *Q* genotypes, 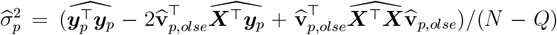 where 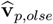is the estimate in Equation (4) with *κ*_*p*_ = *P* . Letting 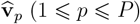 be the estimates with the optimally chosen tuning parameters, the final estimate of **V** is 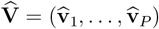

#### 2.2.2. Estimation of super-DAG by the peeling algorithm

Given 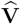, the peeling algorithm (Li et al., 2023) can be used to construct a super-DAG with phenotype edge set *ℰ* ^+^ (a superset of *ℰ* ) and interventional edge set ℐ^+^ (a superset of ℐ). The key idea is that the sparse pattern of matrix **V** characterizes the orientations of the relations among the phenotypes. Specifically, it is demonstrated in Li et al. (2023) that v_*qp*_≠ 0 implies that genotype *q* intervenes on phenotype *p* or an ancestor node of phenotype *p* in the DAG. Thus, if v_*qp*_≠ 0 and v_*qi*_ = 0 for *i*≠ *p*, then phenotype *p* is a leaf node in the DAG, that is, there is no directed edge from phenotype *p* to the others. On this basis, we can sequentially identify and remove (i.e., peel) the leaf node in the DAG, and construct supersets *ℰ* ^+^and ℐ ^+^.

Since the peeling algorithm solely depends on 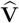, no modification is needed to extend the existing method to accommodate summary-level data.

#### 2.2.3. Estimation of U and W

The peeling algorithm yields supersets *ℰ* ^+^ ⊇ *ℰ* and ℐ^+^ ⊇ ℐ. To remove the extra edges in *ℰ* ^+^ and ℐ^+^, we consider fitting **U** and **W** within a restricted model defined by *ℰ* ^+^ and ℐ^+^.

From Equation (1), for phenotype *p*, we have

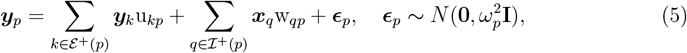

where *ℰ* ^+^(*p*) = *{k* : (*k, p*) ∈ *ℰ* ^+^} and ℐ^+^(*p*) = *{q* : (*q, p*) ∈ ℐ^+^}. As in Section 2.2.1, we replace the corresponding quantities with the summary-level data estimates and fit the TLP regression based on Equation (5),

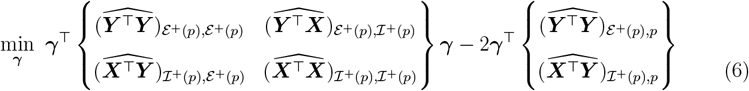

where 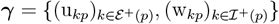 is the parameter vector and . We fix *J* (***γ***, *τ*_*p*) =∑*k*min(|*γk*_|*/τ*_*p*_, 1) . We fix 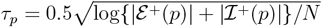 and the tuning parameter *κ*_*p*_ ∈ *{*1, *…*, |*ℰ* ^+^ (*p*)| +|ℐ ^+^ (*p*)|} are selected by pseudo-BIC as described in Section 2.2.1. The estimated 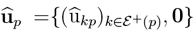 and 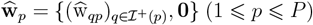 are aggregated to form the the final estimate 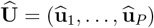 and 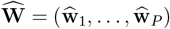.

Due to penalization, we recommend following the common practice to standardize the variables so that the phenotypes and genotypes are on a comparable scale, which is straight-forward to do as 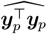 and 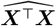 are obtained. Moreover, if only **U** is of interest, penalization of **W** is optional.

### 2.3. Likelihood-based inference for a DAG

We extend the likelihood ratio inference (Li et al., 2023) to quantify the uncertainty of the network structures. As in Li et al. (2023), we consider two types of hypothesis testing.

- **Testing of multiple directed relations**. The null hypothesis *H*_0_ : u_*kj*_ = 0 for each (*k, j*) ∈ *ℋ* and alternative hypothesis *H*_*a*_ : u_*kj*_≠ 0 for some (*k, j*) ∈ *ℋ*. Rejecting *H*_0_ indicates evidence for the presence of some hypothesized relationships in the network.
- **Testing of a directed pathway**. The null hypothesis *H*_0_ : u_*kj*_ = 0 for some (*k, j*) ∈ *ℋ* and alternative hypothesis *H*_*a*_ : u_*kj*_≠ 0 for each (*k, j*) ∈ *ℋ*. Rejecting *H*_0_ indicates evidence for the presence of the entire directed pathway in the phenotype network.

The procedure for testing multiple directed relations comprises five steps.

(T1) Estimate **V** and use the peeling algorithm to obtain *ℰ* ^+^ and ℐ^+^ as in Section 2.2.

(T2) Identify the set of nondegenerate edges 𝒟 (Li et al., 2023), which contains 𝒟_*p*_, the nondegenerate edges pointed to phenotype *p*.

(T3) Estimate the parameters **U** and **W** under *H*_0_ and *H*_*a*_, respectively. Specifically, denote by 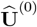 and 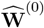 the estimates under *H*_0_. Then 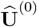 and 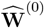 are computed as in the regression (6) with an additional constraint that u_*kj*_ = 0 for (*k, j*) ∈ *ℋ*. Let 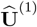 and 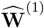 be the estimates under *H*_*a*_. Then 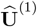 and 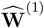 are computed from the restricted models (1 ⩽ *p* ⩽ *P* ), 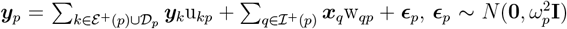 via regression (6), where the penalties become 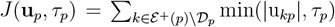 and 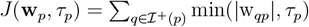

(T4) Compute 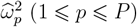 from the residual sum of squares of 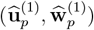

(T5) Compute the test statistic 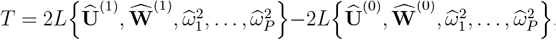 where *L* is the log-likelihood of the model (Equation (1)). By Li et al. (2023), *T* is approximately chi-squared distributed with degrees of freedom |𝒟| when the size | 𝒟| is less than 50; 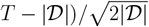 is approximately standard normal when | 𝒟| exceeds 50. Thus, the p-value is calculated as 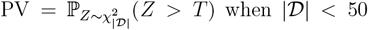 when | 𝒟| < 50 and 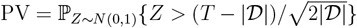 | 𝒟| ⩾ 50.

The procedure for testing a directed pathway is similar, with minor modifications.

(P1) Estimate **V** as in Step (T1).

(P2) First, we decompose *H*_0_ into each nongenerate edge 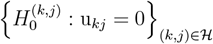 For each 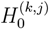 implement Steps (T2)–(T5) above to obtain the corresponding p-value PV_(*k,j*)_. The final p-value is computed as the maximum of the p-values for the sub-hypotheses, PV = max {PV(*k,j*) : (*k, j*) ∈ *ℋ*}.

Of note, testing a directed pathway concerns a composite (null) hypothesis. Fixing 0 *< α <* 1, we have lim_*n→∞*_ sup {ℙ_***θ***_ (PV ⩽ *α*) : ***θ*** = (**U, W**) satisfies *H*0 }= *α* (Li et al., 2023). In other words, the test asymptotically achieves exactly the *α* significance level for the composite null hypothesis.

## 3. Inferring cardiovascular-related protein-protein interaction network

The role of cardiovascular diseases has been recognized as an important etiologic hallmark of AD (de Bruijn and Ikram, 2014). There are different hypotheses on the various mechanisms underlying the association between AD and cardiovascular diseases (Tini et al., 2020). In this real data application, we constructed a directed PPI network of some cardiovascular-related proteins based on a GWAS of 83 plasma protein biomarkers. We further connected the PPI network to AD through MR analyses.

### 3.1. GWAS summary statistics for cardiovascular-related proteins

The GWAS summary statistics on 83 cardiovascular-related proteins which came from Wald tests for the association between each SNP and the standardized residuals among 3394 European individuals by Folkersen et al. (2017) were used. Five proteins were excluded from the analysis as their corresponding protein-encoding genes are located on the sex chromosome. The summary statistics were first processed to remove (a) indels, (b) SNPs located within one base pair of an indel, (c) SNPs with imputation quality score INFO ⩽ 0.8, and (d) SNPs with MAF ⩽ 0.05. We then used the following steps to select putative IVs for the proteins:

- SNPs were clumped at an *r*^2^ value of 0.01 using 3000 uncorrelated individuals (individuals with kinship coefficients less than 0.084) from UK Biobank of European ancestry as the reference panel such that SNPs were independent of each other for each protein (Bycroft et al., 2018);
- Only the SNPs in the clumped data files located within *±*1MB of each protein-encoding genes were considered. In general, cis-regulatory changes will be less pleiotropic (Signor and Nuzhdin, 2018) and thus these SNPs located close to the genes are more likely to be valid IVs due to the exclusion assumption (Swerdlow et al., 2016; Hemani et al., 2018; Li et al., 2023) (i.e., an IV only directly intervenes on one primary variable);
- To ensure the relevance assumption (Li et al., 2023) was satisfied (i.e., IV intervenes on at least one primary variable), we only selected SNPs whose p-values were below the GWAS significance threshold (5 *×* 10^*−*8^). This filtering process led to a total number of 33 SNPs and 23 proteins with at least one putative IV in the final network analysis.

The genetic correlation matrix for the included IVs was estimated based on the same reference panel used in clumping. We calculated the empirical correlation of each pair of proteins as the correlation coefficient of the z-scores of the null SNPs, i.e., all autosomal SNPs with MAF ⩾ 0.05, INFO ⩾ 0.8, and GWAS p-values ⩾ 0.05 for both proteins. The number of null SNPs for each pair of proteins ranged from 1,191,204 to 1,223,357. All preparation of the reference panel and GWAS data for both the DAG estimation and MR analysis was done using PLINK version 1.9 (Purcell et al., 2007).

### 3.2. GWAS summary statistics for AD

We explored the relationship between each of the 23 proteins in Folkersen et al. (2017) and AD. We used the summary statistics of the GWAS for AD from a most recent study totaling 111,326 clinically diagnosed or “proxy” AD cases and 677,663 controls (Bellenguez et al., 2022). We removed SNPs with MAF < 0.05, SNPs not included in the GWAS of Folkersen et al. (2017), and clumped SNPs at *r*^2^ = 0.01. Among the remaining SNPs, we selected IVs only with GWAS p-value < 5 *×* 10^*−*8^ in the MR analyses.

### 3.3. Results

We constructed a DAG of the 23 proteins as described in Section 2.5.1. As Folkersen et al. (2017) shared MAF in the summary statistics, we compared them with those in UK Biobank, the reference panel for clumping and estimating genetic correlation matrix. The absolute difference of the MAF of all IVs ranged from 0.001 to 0.055 with a mean 0.02, while the correlation was 0.99 (Table S1). We further performed MR analysis on each protein to evaluate their relationship with AD using the TwoSampleMR package (Hemani et al., 2018). We used Egger’s test of intercept for examining the exclusion assumption: If a protein had a p-value of the Egger’s test of intercept *>* 0.05*/*23, there was no evidence against no direct/pleiotropic effects, and we’d go with the more powerful MR-IVW method; otherwise, we used MR-Egger (to allow pleiotropic effects of IVs). In any case, we used the p-value cut-off *<* 0.05*/*23 to declare statistical significance. Table S2 contains a complete list of MR results. The protein IL18, which showed marginal significance in both Egger’s test of intercept (p-value = 0.06) and MR-Egger (p-value = 0.07), was a parent node for several proteins related to AD, including ADM, IL1RL1, CTSD, CXCL6, and CXCL16. We further performed the likelihood ratio test on each edge. Edges with p-value *<* 0.05*/*(23 *×* 22 *−* 56) were considered as significant and were in solid line in Figure 2. The number of tests in the Bonferonni correction is bounded by the sum of possible edges among all the nodes minus the total number of edges after the peeling algorithm, which is justified in Supplementary Material S2.1. Each edge from IL18 to the five AD-associated proteins was highly significant in the likelihood ratio test, thus suggesting that simultaneous testing of the pathway from IL18 to the five genes would be significant. Previous studies detected increased levels of pro-inflammatory IL18 in both cardiovascular diseases and in brain regions of AD patients (Sutinen et al., 2014). IL18 is known to increase the level of Cdk5 and GSK-3*β*, which are involved in Tau hyperphosphorylation, and the inhibition of Cdk5 was known to improve AD subject’s condition (Calabrò et al., 2021). Our work suggests a possible regulatory role of IL18 on multiple AD-associated proteins. According to OpenTargets.org (Ochoa, 2021) for current pharmaceuticals either approved or in development with IL18, this protein is currently a target of an antibody drug to treat diabetes mellitus and a few other conditions; diabetes has long been linked to AD with epidemiological and biological evidence (Barbagallo and Dominguez, 2014). Lastly, we provide a Shiny application that allows users to test any selected proteins in this cardiovascular-related PPI network.

**Figure 1.**
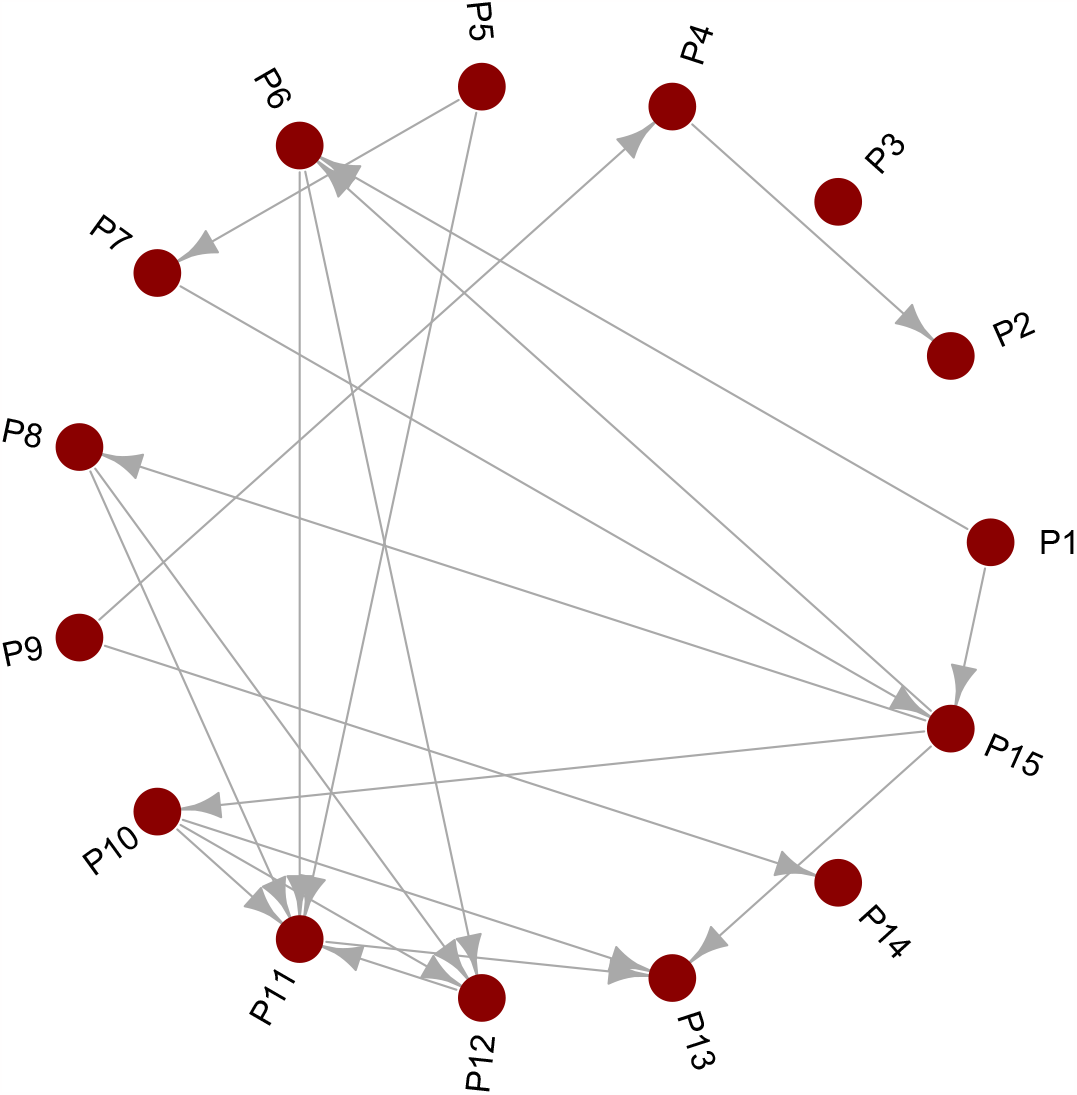
True DAG for the simulation study with 15 phenotypes

**Figure 2.**
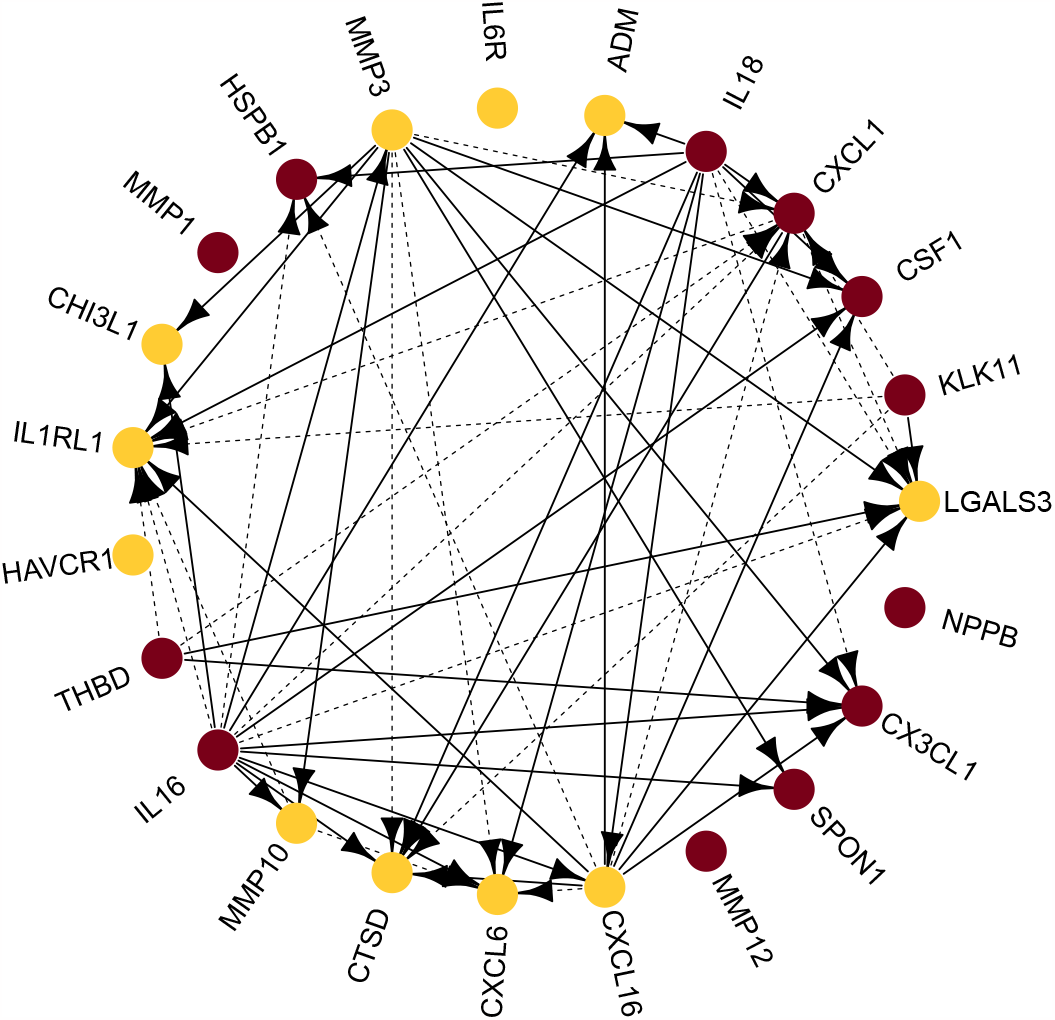
Estimated DAG for 23 proteins based on the GWAS summary statistics of Folkersen et al. (2017); Proteins significantly associated with AD in MR analysis are colored gold. A solid line represents an edge that is statistically significant by the likelihood ratio test whereas a dashed line represents an edge that is not significant.

## 4 Simulation studies

### 4.1. Simulation settings

We simulated the data assuming a fixed **U, W**, standardized genotype matrix ***X***, and sampled each row of ***ϵ*** independently from 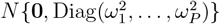 Then we generated ***Y*** from equation: ***Y*** = **V**^*T*^***X*** + ***ϵ***(**I** *−* **U**)^*−*1^. Without loss of generality, no intercept was modeled, i.e., ***Y*** was centered at mean 0. The values of 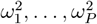 and **W** are provided in Supplemental Material S1.1, where the structure of the relationship of ***Y*** followed a DAG of 15 nodes/phenotypes (Figure 1). The effect sizes of the non-zero components of **U** ranged from 0.002 to 1.16 with a median of 0.06. All phenotypes had at least one valid IV. Twenty-six SNPs were included in the model, with their effect sizes ranging from *−*2.2 to 2.5 with a median of *−*0.11. Two SNPs violated the relevance assumption, while the rest were valid IVs. We also varied the effect sizes to be 1*/*3 and 1*/*15 of **U** while keeping **W** fixed.

The standardized genotype matrix ***X*** was obtained from unrelated individuals of European ancestry in UK Biobank (Bycroft et al., 2018). We then calculated the summary statistics using a linear model of each phenotype on each standardized genotype, and inputted the summary statistics into the proposed algorithm. The reference panel was obtained from the UK Biobank European samples, which were not correlated with the simulated samples used to derive the summary statistics. SNPs on chromosome 22 with a Hardy-Weinberg Disequilibrium test p-value *>* 0.0001, missing call rate *<* 0.05, and MAF *>* 0.05 were pruned to have *r*^2^ *<* 0.01. We then randomly selected 26 SNPs for ***X***. Missing values of SNPs were imputed by their mean. Null SNPs were directly simulated to be independent of each other and had no relationship with ***Y***.

We evaluated the performance of both the network construction and statistical inference for the proposed method. To evaluate the performance of network construction, we examined the false positive (TP) and false negative (FN) rates for 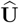 over 200 replications. The sample size of the summary statistics was varied at 3000, 6000, 9000, and 12,000, and the sample size of the reference panel was fixed at 3000. Null SNPs were simulated to have the same sample size as the GWAS summary statistics.

In terms of testing, we examined the empirical Type I error/power of the likelihood ratio tests with increasing sample sizes and varying strength of **U** for the following five scenarios over 1000 replications for the two types of testing.

#### I. Testing one or more directed relations

- A1. Type 1 error: testing one edge when in truth it was null with *H*_0_ : u_1,14_ = 0 vs *H*_1_ : u_1,14_≠ 0 ;
- A2. Power: testing one edge when in truth it was not null with *H*_0_ : u_1,6_ = 0 vs *H*_1_ : u_1,6_≠ 0;
- A3. Power: testing two edges together when in truth both were not null with *H*_0_ : u_7,15_ = u_1,6_ = 0 vs *H*_1_ : u_7,15_ ≠ 0 or u_1,6_≠ 0;

#### II. Testing of a directed pathway

- B1. Type 1 error: testing whether at least one edge was not null when in truth only one was not null with *H*_0_ : u_1,14_ = 0 or u_6,12_ = 0 vs *H*_1_ : u_1,14_ ≠ 0 and u_6,12_≠ 0;
- B2. Power: testing whether at least one edge was not null when in truth both were not null with *H*_0_ : u_1,6_ = 0 or u_6,12_ = 0 vs *H*_1_ : u_1,6_≠ 0 and u_6,12_≠ 0;

The true strengths of the tested edges were: u_1,14_ = 0, u_1,6_ = 0.27, u_7,15_ = 0.44, and u_6,12_ = 1.36.

### 4.2. Simulation results

With the increase of sample size, the constructed networks became closer to the true graph (Figure 3, Numeric values in Figure 3. A are in Table S3 - S6). More specifically, FP of 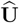 was around 0.05, and FN of 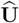 decreased with the increase of sample size when 15000 null SNPs were used to estimate the 15 *×* 15 matrix of ***Y*** ^*T*^***Y*** . Unsurprisingly, the performance of using 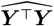 estimated by 15,000 null SNPs (denoted as **U**^*′*^ in Figure 3) was slightly worse than that of using ***Y*** ^*T*^***Y*** (denoted as **U** in Figure 3). In addition, the FP of 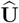 clearly decreased with the increase of effect sizes (from **U***/*15 to **U***/*3 to **U**). To check the validity of our method, we compared pBIC estimated from summary statistics with BIC estimated from the individual-level data for the same set of penalized regression coefficients. We found the two sets of values were highly concordant, and the results from one iteration were plotted in Figure S2.

**Figure 3.**
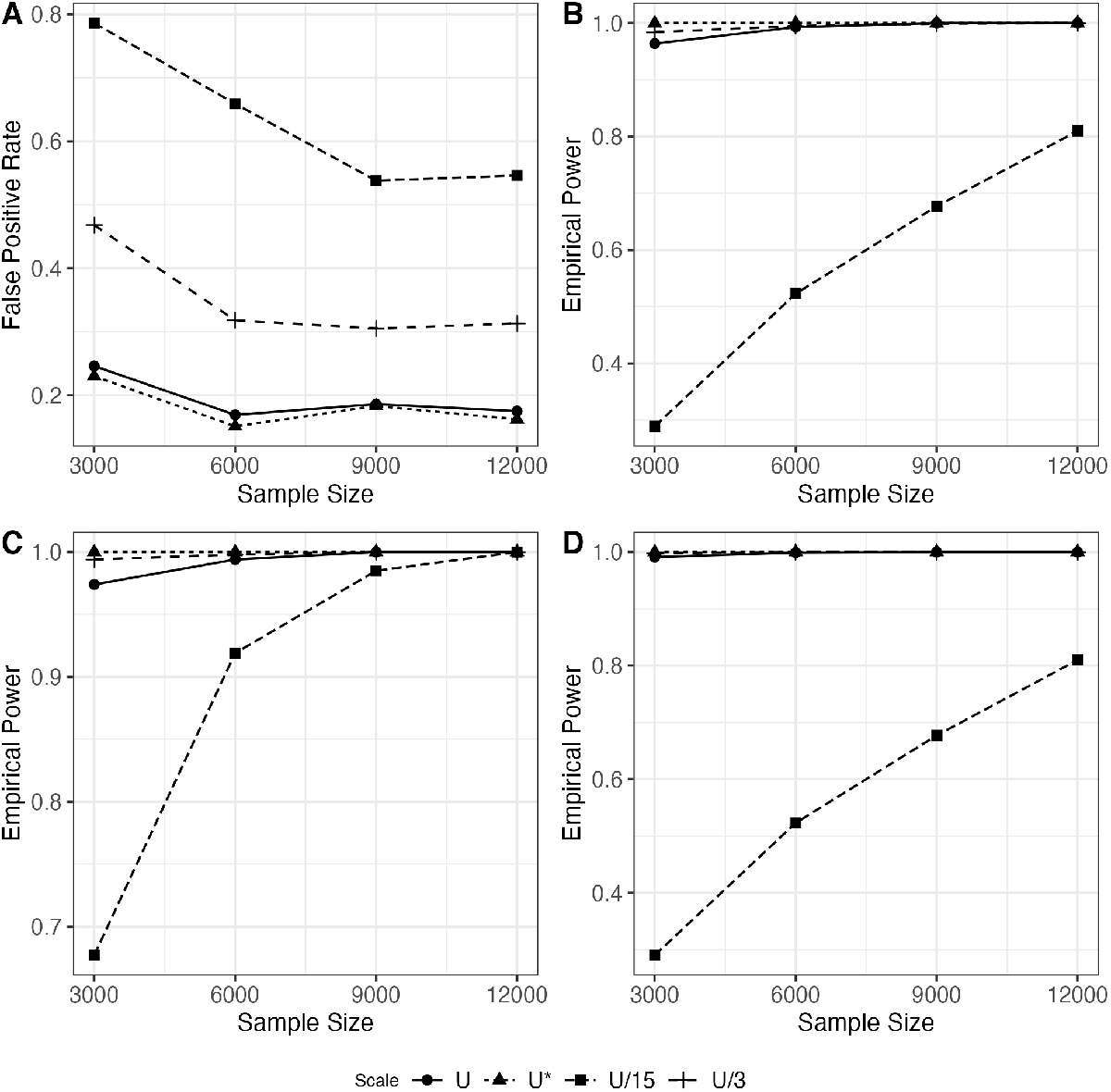
Performance of network construction (A) and likelihood ratio tests (B-D) in simulation with varying sample sizes and true effect sizes at **U, U***/*3, and **U***/*15. Figures B, C, and D represent scenarios A2, A3, and B2 respectively.

In terms of testing, we observed well-controlled Type I error rates for scenarios A1 and B1 (Numeric values present in Figure 3, B-D are in Table S7 - S10), using ***Y*** ^*T*^***Y*** . We note that the empirical Type I error rates might become conservative when ***Y*** ^*T*^***Y*** is replaced by its estimate 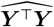 derived from 15000 null SNPs (Table S10). In real data analysis of GWAS summary level data, typically a much larger number of null SNPs and various other strategies can be used to better estimate ***Y*** ^*T*^***Y*** (Bulik-Sullivan et al., 2015; Kim et al., 2015; Li et al., 2021); however, the investigation in this direction is out of our scope. Furthermore, empirical power was high for scenarios A2, A3, and B2 and increased with sample size and effect size. The empirically power of jointly testing two edges u_1,6_ and u_7,15_ (scenario A3) was larger than testing one edge u_1,6_ alone (scenario A2).

## 5. Discussion

In this paper, we present a method to estimate an interventional DAG of phenotypes utilizing linear structural equation models, applicable to GWAS summary statistics in the absence of individual-level data. We demonstrated satisfactory performance in terms of the FP and FN rates in network construction and high empirical power and of well-controlled Type I error rates of the likelihood ratio tests. We applied this method to a large-scale proteomic GWAS summary dataset to obtain an estimated DAG of 23 cardiovascular-related proteins and further illustrated the effects of these proteins on AD by MR analysis. These results can be useful in understanding the disease etiology, drug repurposing, and other applications for AD.

We note that the choice of a proper reference panel is just as important for our method as many other summary-statistics-based methods (Deng and Pan, 2018; Chen et al., 2021; Privéet al., 2022). When constructing the cardiovascular-related PPI network, we used an ancestry-matched reference panel of uncorrelated individuals from UK Biobank with a sample size of 3000, which is close to the sample size of the GWAS of cardiovascular proteins. Furthermore, we clumped SNPs around the cis-region of the gene and only used the genome-wide significant SNPs as IVs. This step not only aimed at selecting at least one valid IV for each protein, but also achieving better estimation of the genetic correlation matrix for the IVs as the number of total IVs became much smaller than the sample size of the reference panel, i.e. *Q ≪ N*_*r*_. Our analysis was constrained to a single ancestry group and unrelated samples. As the collection of multi-ancestry and related samples increases, it will be of significant research interest to establish networks among these populations, a challenge we anticipate addressing in our future work.

In recent years, a large amount of summary-level data has become widely accessible. On the molecular level, many studies published their summary statistics for SNP-molecular phenotype associations. For example, variant-gene associations in 49 tissues can be directly downloaded from GTExPortal (Consortium, 2020). Beyond the molecular phenotypes, UK Biobank alone provides GWAS summary statistics on more than 7000 traits, including but not limited to cognitive functions, early life factors, health and medical history, and physical measurement (Bycroft et al., 2018). Our proposed method provides a computational and analytical tool to explore the relationships among multiple phenotype variables by taking advantage of rapid advances in GWAS and other association mappings.

## Acknowledgement

R. Zilinskas and C. Li have contributed equally to this work. The authors would like to thank the associate editor and the reviewer for their valuable comments. This research was supported by NIH grants U01 AG073079, R01 AG065636, R01 AG069895, RF1 AG067924, R01 HL116720, and R01 GM126002 and by the Minnesota Supercomputing Institute at the University of Minnesota. T. Yang would like to further acknowledge the Children’s Cancer Research Funds and the St. Baldrick’s Foundation Scholar Award.

## Supplementary Materials

Web Appendices, Supplementary Tables S1–S10, and Figure S1 referenced in Sections 3.3 and 4.2 are available with this paper in the Supplementary Materials. Data and code (sumDAG algorithm, Shiny app, and the real data analysis code) can also be found on https://github.com/chunlinli/sumdag.

## Data availability statement

We downloaded the summary level GWAS data in Section 3.1 from https://zenodo.org/record/264128/. The algorithm for the proposed work is packaged in R, available at https://github.com/chunlinli/sumdag, along with code used for the simulation studies and real data application. The processed summary-level GWAS data that were used as input for the algorithm for the real data application are also included on GitHub.

## Notes

### Competing Interest Statement

The authors have declared no competing interest.

### Summary of Updates

We have made clarifications explaining our methods and terms across the manuscript. We further emphasized the importance of the choice of reference panel for our method in the Discussion section. For our real data analysis, we applied more restrictive criteria to select the instrumental variables for the cardiovascular-related proteins and changed the reference panel from 1000G to UK Biobank, which has a similar sample size of the GWAS summary statistics for pruning and estimating the genetic correlation matrix. We found that UK Biobank generates a more precise estimation of MAF, especially for these instrumental variables with relatively lower MAF than using 1000G. Given these changes, we have observed some alterations in the constructed network (as illustrated in Figure 2); Nevertheless, IL18 has remained to be a protein of potential interest for Alzheimer's Disease. To enhance the reproducibility of our findings, we present an R package, updated code for both simulation and real data applications, and a Shiny application at github.com/chunlinli/sumdag. The GitHub page includes the GWAS summary statistics files, so that readers can have a better understanding of the input files of our algorithm and use them to replicate the real data application. The Shiny application allows users to test any selected proteins, providing more flexibility for those interested in understanding the cardiovascular-related protein-protein interaction network. All of the current content on the GitHub page has also been provided as Supporting Information as suggested.

