## Supplementary Materials for "Inferring a directed acyclic graph of phenotypes from GWAS summary statistics"

### Supplementary Materials for "Inferring a directed acylic graph of phenotypes from GWAS summary statistics" by Zilinskas, Li, Shen, Pan and Yang

### S1 Simulation Studies

#### S1.1 True assumed DAGs for simulation studies

$\boldsymbol{U}$ rounded to 2 decimal points was set as:

| | $P1$ | $P2$ | $P3$ | $P4$ | $P5$ | $P6$ | $P7$ | $P8$ | $P9$ | $P10$ | $P11$ | $P12$ | $P13$ | $P14$ | $P15$ |
|---|---|---|---|---|---|---|---|---|---|---|---|---|---|---|---|
| $P1$ | 0 | 0 | 0 | 0 | 0 | 0.27 | 0 | 0 | 0 | 0 | 0 | 0 | 0 | 0 | 1.87 |
| $P2$ | 0 | 0 | 0 | 0 | 0 | 0 | 0 | 0 | 0 | 0 | 0 | 0 | 0 | 0 | 0 |
| $P3$ | 0 | 0 | 0 | 0 | 0 | 0 | 0 | 0 | 0 | 0 | 0 | 0 | 0 | 0 | 0 |
| $P4$ | 0 | 0.94 | 0 | 0 | 0 | 0 | 0 | 0 | 0 | 0 | 0 | 0 | 0 | 0 | 0 |
| $P5$ | 0 | 0 | 0 | 0 | 0 | 0 | 0.95 | 0 | 0 | 0 | 1.18 | 0 | 0 | 0 | 0 |
| $P6$ | 0 | 0 | 0 | 0 | 0 | 0 | 0 | 0 | 0 | 0 | 1.13 | 1.36 | 0 | 0 | 0 |
| $P7$ | 0 | 0 | 0 | 0 | 0 | 0 | 0 | 0 | 0 | 0 | 0 | 0 | 0 | 0 | 0.44 |
| $P8$ | 0 | 0 | 0 | 0 | 0 | 0 | 0 | 0 | 0 | 0 | 0.46 | 0.02 | 0 | 0 | 0 |
| $P9$ | 0 | 0 | 0 | 1.13 | 0 | 0 | 0 | 0 | 0 | 0 | 0 | 0 | 0 | 1.28 | 0 |
| $P10$ | 0 | 0 | 0 | 0 | 0 | 0 | 0 | 0 | 0 | 0 | 1.51 | 1.32 | 0.10 | 0 | 0 |
| $P11$ | 0 | 0 | 0 | 0 | 0 | 0 | 0 | 0 | 0 | 0 | 0 | 0 | 0.24 | 0 | 0 |
| $P13$ | 0 | 0 | 0 | 0 | 0 | 0 | 0 | 0 | 0 | 0 | 0 | 0 | 0 | 0 | 0 |
| $P14$ | 0 | 0 | 0 | 0 | 0 | 0 | 0 | 0 | 0 | 0 | 0 | 0 | 0 | 0 | 0 |
| $P15$ | 0 | 0 | 0 | 0 | 0 | 0.94 | 0 | 2.37 | 0 | 1.20 | 0 | 0 | 0.38 | 0 | 0 |

$\boldsymbol{W}$ matrix round to the 2 decimal points was set as:

| | $P1$ | $P2$ | $P3$ | $P4$ | $P5$ | $P6$ | $P7$ | $P8$ | $P9$ | $P10$ | $P11$ | $P12$ | $P13$ | $P14$ | $P15$ |
|---|---|---|---|---|---|---|---|---|---|---|---|---|---|---|---|
| $SNP1$ | 0 | 0 | 0 | 0 | 0 | 0.66 | 0 | 0 | 0 | 0 | 0 | 0 | 0 | 0 | 0 |
| $SNP2$ | 0 | 0 | 0 | 0 | 0 | 0 | 0 | −0.44 | 0 | 0 | 0 | 0 | 0 | 0 | 0 |
| $SNP3$ | 0 | 0 | 0 | 0 | 0 | 0 | 0 | 0 | 0 | 2.47 | 0 | 0 | 0 | 0 | 0 |
| $SNP4$ | 0 | 0 | 0 | 0 | 0 | 0 | 0 | 0 | 0 | 0 | 0 | −2.26 | 0 | 0 | 0 |
| $SNP5$ | 0 | 0 | 0 | 0 | 0 | 0 | 0 | 0 | 0 | 2.34 | 0 | 0 | 0 | 0 | 0 |
| $SNP6$ | 0 | 0 | 0 | 0 | 0 | 0 | 0 | 0 | 0 | −1.08 | 0 | 0 | 0 | 0 | 0 |
| $SNP7$ | 0 | 0 | 0 | 0 | 0 | 0 | 0 | 0 | 0 | −1.16 | 0 | 0 | 0 | 0 | 0 |
| $SNP8$ | 0 | 0 | 0 | 0 | 0 | 0 | 0 | 0 | 0 | 0 | −1.73 | 0 | 0 | 0 | 0 |
| $SNP9$ | 0 | 0 | 0 | 0 | 0 | 0 | 0 | 0 | 0 | 0 | 0 | 0 | 0 | 0 | −1.86 |
| $SNP10$ | 0 | 0 | 0 | 0 | 0 | 0 | 0 | 0 | 0 | 0 | 0 | 0 | 0 | 0 | 0 |
| $SNP11$ | 0 | 0 | 0 | 0 | 0 | 0 | 0 | 0 | 0 | 0 | 0 | 0 | 0.34 | 0 | 0 |
| $SNP12$ | 0 | 0 | 0 | 0 | 0 | 0 | 0 | 0 | 0 | 0 | 0 | 0 | 0 | 0 | 0 |
| $SNP13$ | 0 | 0 | 0 | 0 | 0 | 0 | 0 | 0 | 0 | 0 | 0 | 0 | 0 | 0.66 | 0 |
| $SNP14$ | 0 | 0 | −1.19 | 0 | 0 | 0 | 0 | 0 | 0 | 0 | 0 | 0 | 0 | 0 | 0 |
| $SNP15$ | 0 | 0 | 0 | 0 | −0.88 | 0 | 0 | 0 | 0 | 0 | 0 | 0 | 0 | 0 | 0 |
| $SNP16$ | 0 | 0 | 0 | 0 | 0 | 0 | 0 | 0 | 0 | 0 | 0 | 0 | 0 | 0 | −0.87 |
| $SNP17$ | 0.37 | 0 | 0 | 0 | 0 | 0 | 0 | 0 | 0 | 0 | 0 | 0 | 0 | 0 | 0 |
| $SNP18$ | 0 | 0 | 0 | 0 | 0 | 0 | 0 | 0 | 0 | 0 | 0 | 0 | 0 | 0 | −1.02 |
| $SNP19$ | 0 | 0 | 0 | 0 | 0 | 0 | 1.56 | 0 | 0 | 0 | 0 | 0 | 0 | 0 | 0 |
| $SNP20$ | 0 | 0 | 0 | 0 | 0 | 0 | 0 | 0 | 0 | 0 | 0 | 0 | −1.25 | 0 | 0 |
| $SNP21$ | −1.23 | 0 | 0 | 0 | 0 | 0 | 0 | 0 | 0 | 0 | 0 | 0 | 0 | 0 | 0 |
| $SNP22$ | 0 | 0 | 0 | 0 | 0.66 | 0 | 0 | 0 | 0 | 0 | 0 | 0 | 0 | 0 | 0 |
| $SNP23$ | 0 | 0 | 0 | 0 | 0 | 0 | 0 | 0 | 0.44 | 0 | 0 | 0 | 0 | 0 | 0 |
| $SNP24$ | 0 | 0 | 0 | 0 | 0 | 0 | 0 | 0 | 0 | 0.24 | 0 | 0 | 0 | 0 | 0 |
| $SNP25$ | 0 | 0 | 0 | 0.35 | 0 | 0 | 0 | 0 | 0 | 0 | 0 | 0 | 0 | 0 | 0 |
| $SNP26$ | 0 | 0.28 | 0 | 0 | 0 | 0 | 0 | 0 | 0 | 0 | 0 | 0 | 0 | 0 | 0 |

$\omega_1^2, \ldots, \omega_P^2$ were set as 0.64, 0.56, 0.73, 0.71, 0.90, 0.66, 0.64, 0.57, 0.78, 0.63, 0.76, 0.47, 0.83, 0.60, and 0.58.

### S2 Real data application results

#### S2.1 Adjustment for multiple testing

Consider testing of directed relations $H_0^{(k,j)}$ : $\mathrm{u}_{kj} = 0$ against its alternative; $1 \leq k \neq j \leq P$. To control the Type-I error rate,

$$\begin{aligned}
& P\Big(\text{some } H_0^{(k,j)} \text{ is rejected} \mid \mathcal{G}\Big) \\
\leq & \sum_{(k,j)\notin\mathcal{E}} P\Big(H_0^{(k,j)} \text{ is rejected}\Big) \\
= & \sum_{(k,j)\notin\mathcal{E}:\{(k,j)\}\cup\mathcal{E}^+ \text{ acyclic}} P\Big(H_0^{(k,j)} \text{ is rejected}\Big) + \underbrace{\sum_{(k,j):\{(k,j)\}\cup\mathcal{E}^+ \text{ cyclic}} P\Big(H_0^{(k,j)} \text{ is rejected}\Big)}_{\text{degenerate cases, no rejection}} \\
\approx & \sum_{(k,j):\{(k,j)\}\cup\mathcal{E}^+ \text{ acyclic}} P\Big(H_0^{(k,j)} \text{ is rejected}\Big).
\end{aligned}$$

The number of test to adjust is $p^2 - p - |\mathcal{E}^+|$.

#### S2.2 MAF for IVs in GWAS and reference panel

Table S1: MAF of 33 IVs in GWAS summary statistics of Folkerson et al and reference panel

| SNP | maf(ukbb) | maf(Folkersen) |
|---|---:|---:|
| rs1048328 | 0.93 | 0.91 |
| rs10920579 | 0.80 | 0.79 |
| rs11084045 | 0.94 | 0.95 |
| rs11225399 | 0.73 | 0.77 |
| rs113299890 | 0.92 | 0.92 |
| rs116187470 | 0.87 | 0.82 |
| rs12127600 | 0.60 | 0.61 |
| rs1420101 | 0.61 | 0.63 |
| rs1580006 | 0.51 | 0.54 |
| rs16850178 | 0.94 | 0.95 |
| rs17610659 | 0.47 | 0.48 |
| rs17875523 | 0.90 | 0.94 |
| rs1969539 | 0.51 | 0.50 |
| rs198379 | 0.56 | 0.60 |
| rs2241132 | 0.82 | 0.85 |
| rs2868371 | 0.77 | 0.77 |
| rs2886922 | 0.57 | 0.60 |
| rs3176123 | 0.80 | 0.81 |
| rs34393565 | 0.84 | 0.85 |
| rs35045092 | 0.77 | 0.78 |
| rs4129267 | 0.61 | 0.64 |
| rs4235719 | 0.62 | 0.61 |
| rs471994 | 0.63 | 0.65 |
| rs481814 | 0.96 | 0.94 |
| rs549596 | 0.54 | 0.59 |
| rs56105155 | 0.89 | 0.89 |
| rs61868977 | 0.54 | 0.56 |
| rs6555820 | 0.52 | 0.52 |
| rs670211 | 0.45 | 0.41 |
| rs7533952 | 0.57 | 0.57 |
| rs75649625 | 0.72 | 0.76 |
| rs7943617 | 0.74 | 0.77 |
| rs7946057 | 0.49 | 0.53 |

#### S2.3 MR results

Table S2: MR analysis results for the 23 proteins and AD, nIV is the number of SNPs included as IV

| Gene | nIV | MR-Egger | Weighted Median | IVW | Egger's test of intercept |
|---|---|---|---|---|---|
| ADM | 407 | 6.8E-01 | 9.1E-28 | 4.9E-26 | 3.1E-01 |
| LGALS3 | 840 | 2.5E-55 | 1.1E-177 | 2.5E-81 | 2.5E-11 |
| KLK11 | 31 | 7.1E-01 | 7.2E-01 | 8.1E-01 | 7.7E-01 |
| CSF1 | 7 | 1.5E-01 | 6.0E-01 | 6.9E-01 | 1.6E-01 |
| CXCL1 | 34 | 1.9E-01 | 8.6E-01 | 3.6E-01 | 2.2E-01 |
| IL18 | 110 | 7.2E-02 | 4.5E-01 | 9.5E-01 | 6.4E-02 |
| IL6R | 430 | 1.2E-03 | 1.3E-29 | 4.7E-46 | 1.5E-03 |
| MMP3 | 339 | 4.1E-01 | 5.7E-04 | 4.8E-07 | 4.1E-01 |
| HSPB1 | 48 | 7.6E-01 | 1.5E-01 | 4.6E-02 | 9.8E-01 |
| MMP1 | 119 | 6.1E-01 | 1.8E-03 | 1.1E-02 | 2.9E-01 |
| CHI3L1 | 146 | 5.5E-01 | 1.9E-06 | 2.9E-12 | 2.0E-01 |
| IL1RL1 | 661 | 1.3E-05 | 7.9E-45 | 7.1E-47 | 6.2E-01 |
| HAVCR1 | 222 | 1.9E-03 | 1.0E+00 | 1.6E-07 | 9.3E-06 |
| THBD | 60 | 4.9E-01 | 8.6E-01 | 6.6E-02 | 1.4E-01 |
| IL16 | 28 | 7.7E-01 | 3.5E-10 | 2.5E-03 | 3.5E-01 |
| MMP10 | 161 | 3.5E-01 | 7.4E-12 | 3.4E-16 | 1.1E-02 |
| CTSD | 137 | 3.4E-04 | 1.6E-05 | 1.5E-01 | 9.2E-04 |
| CXCL6 | 81 | 6.2E-01 | 2.3E-07 | 2.0E-09 | 1.5E-01 |
| CXCL16 | 76 | 8.5E-01 | 1.0E-05 | 1.4E-05 | 4.5E-01 |
| MMP12 | 386 | 1.1E-01 | 4.9E-01 | 6.0E-01 | 1.3E-01 |
| SPON1 | 154 | 8.6E-01 | 3.9E-01 | 9.2E-02 | 9.9E-01 |
| CX3CL1 | 19 | 2.9E-01 | 2.4E-01 | 3.9E-01 | 3.2E-01 |
| NPPB | 116 | 1.6E-01 | 2.1E-36 | 5.1E-22 | 5.2E-03 |

### S3 More simulation results

#### S3.1 pseudo BIC

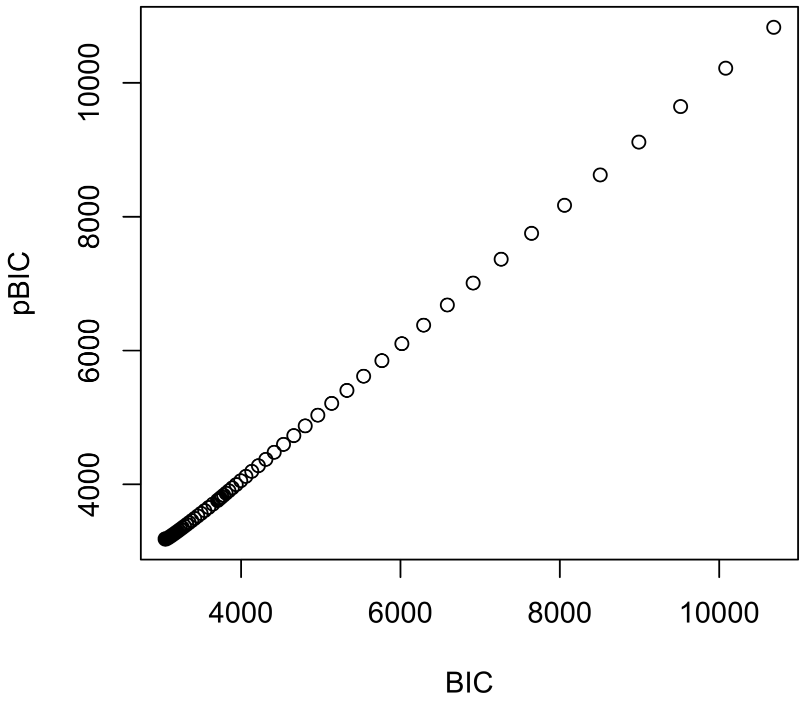


Figure S1: Comparison of BIC and pBIC under 1 simulation iteration (sample size = 3000)

#### S3.2 Network Construction

Table S4 and Table S5 evaluate the summary statistics with $\boldsymbol{Y^TY}$ estimated by empirical Y where Table S4 evaluates $\boldsymbol{U}$ with 1/3 of the values of $\boldsymbol{U}$ in Section S1.1 and Table S5 evaluates $\boldsymbol{U}$ with 1/15 of the values of $\boldsymbol{U}$ in Section S1.1.

Table S3: Network construction using summary statistics with $\boldsymbol{U}$

| N | FP($\hat{\boldsymbol{V}}$) | FN($\hat{\boldsymbol{V}}$) | FP($\hat{\boldsymbol{U}}$) | FN($\hat{\boldsymbol{U}}$) |
|---|---|---|---|---|
| 3000 | 0.091 | 0.087 | 0.040 | 0.246 |
| 6000 | 0.073 | 0.061 | 0.029 | 0.169 |
| 9000 | 0.073 | 0.068 | 0.038 | 0.186 |
| 12000 | 0.076 | 0.063 | 0.036 | 0.175 |

Table S4: Network construction using summary statistics with $\boldsymbol{U}$ and $\boldsymbol{Y^TY}$ estimated by 15,000 null SNPs

| N | FP($\hat{\boldsymbol{V}}$) | FN($\hat{\boldsymbol{V}}$) | FP($\hat{\boldsymbol{U}}$) | FN($\hat{\boldsymbol{U}}$) |
|---|---|---|---|---|
| 3000 | 0.091 | 0.088 | 0.053 | 0.230 |
| 6000 | 0.073 | 0.059 | 0.058 | 0.151 |
| 9000 | 0.072 | 0.065 | 0.066 | 0.183 |
| 12000 | 0.076 | 0.062 | 0.070 | 0.162 |

Table S5: Network construction using summary statistics with $\boldsymbol{U}/3$

| N | FP($\hat{\boldsymbol{V}}$) | FN($\hat{\boldsymbol{V}}$) | FP($\hat{\boldsymbol{U}}$) | FN($\hat{\boldsymbol{U}}$) |
|---|---|---|---|---|
| 3000 | 0.036 | 0.359 | 0.015 | 0.468 |
| 6000 | 0.033 | 0.292 | 0.022 | 0.318 |
| 9000 | 0.029 | 0.274 | 0.022 | 0.305 |
| 12000 | 0.033 | 0.277 | 0.015 | 0.313 |

Table S6: Network construction using summary statistics with $\boldsymbol{U}/15$

| N | FP($\hat{\boldsymbol{V}}$) | FN($\hat{\boldsymbol{V}}$) | FP($\hat{\boldsymbol{U}}$) | FN($\hat{\boldsymbol{U}}$) |
|---|---|---|---|---|
| 3000 | 0.182 | 0.506 | 0.017 | 0.786 |
| 6000 | 0.214 | 0.344 | 0.025 | 0.659 |
| 9000 | 0.260 | 0.303 | 0.024 | 0.538 |
| 12000 | 0.262 | 0.324 | 0.024 | 0.546 |

#### S3.3 Testing

Table S7: Empirical power and Type I error for the likelihood ratio test (with $\boldsymbol{U}$)

| N | A1 | A2 | A3 | B1 | B2 |
|---|---|---|---|---|---|
| 3000 | 0.055 | 0.964 | 0.974 | 0.068 | 0.991 |
| 6000 | 0.050 | 0.993 | 0.994 | 0.054 | 0.999 |
| 9000 | 0.050 | 1 | 1 | 0.050 | 1 |
| 12000 | 0.060 | 1 | 1 | 0.060 | 1 |

Table S8: Empirical power and Type I error for the likelihood ratio test (with $\boldsymbol{U}/3$)

| N | A1 | A2 | A3 | B1 | B2 |
|---|---|---|---|---|---|
| 3000 | 0.048 | 0.984 | 0.994 | 0.048 | 0.998 |
| 6000 | 0.053 | 0.995 | 0.998 | 0.053 | 1 |
| 9000 | 0.055 | 0.999 | 1 | 0.055 | 1 |
| 12000 | 0.044 | 1 | 1 | 0.044 | 1 |

Table S9: Empirical power and Type I error for the likelihood ratio test (with $\boldsymbol{U}/15$)

| N | A1 | A2 | A3 | B1 | B2 |
|---|---|---|---|---|---|
| 3000 | 0.054 | 0.289 | 0.677 | 0.054 | 0.290 |
| 6000 | 0.049 | 0.523 | 0.919 | 0.049 | 0.523 |
| 9000 | 0.048 | 0.677 | 0.985 | 0.048 | 0.677 |
| 12000 | 0.043 | 0.810 | 1 | 0.043 | 0.810 |

Table S10: Empirical power and Type I error for the likelihood ratio test with (with $\boldsymbol{U}/15$) and $\boldsymbol{Y}^{\boldsymbol{T}}\boldsymbol{Y}$ estimated by 15,000 null SNPs

| N | A1 | A2 | A3 | B1 | B2 |
|---|---|---|---|---|---|
| 3000 | 0.013 | 0.974 | 0.997 | 0.012 | 0.944 |
| 6000 | 0 | 0.874 | 0.952 | 0 | 0.762 |
| 9000 | 0 | 0.971 | 0.997 | 0 | 0.970 |
| 12000 | 0.001 | 0.970 | 0.994 | 0 | 0.965 |
